# Measurement of residual stress in the brain

**DOI:** 10.1101/2025.02.26.637925

**Authors:** Ramin Balouchzadeh, Christopher D. Kroenke, Kara E. Garcia, Philip V. Bayly

## Abstract

Mechanical stress in brain tissue is an important feature of the brain, likely to play key roles in brain development and brain injury. Here, we characterize residual stresses in grey matter and white matter from the deformed shape of tissue cylinders extracted from the brain using a biopsy needle and imaged with high-resolution magnetic resonance imaging (MRI). We use finite element simulations to reconstruct the stress state of tissue sections in the intact brain from images of corresponding sections of the deformed excised tissue. In both adult mouse and Yucatan minipig brains, cortical grey matter exhibited predominantly compressive stresses, while white matter exhibited strongly anisotropic tensile stresses. The direction of maximum tension in white matter generally aligns with axon orientation as observed with diffusion MRI in the minipig. These stress patterns (compressive in cortical grey matter, tensile along axons) are consistent with constrained cortical expansion and tension-induced axonal growth during brain development. These findings offer new insights into the biomechanical factors underlying brain morphogenesis, with implications for understanding neurodevelopmental disorders, brain injuries, and neurosurgical interventions.

## INTRODUCTION

Residual stress, the internal mechanical stress that persists within a structure in the absence of external loads, plays an important role in the mechanical integrity and function of many biological tissues. Residual stress has been extensively studied in arteries, tumors, and skin, where its measurement has provided insights into physiology, including the understanding of tissue elasticity and the role of mechanical factors in maintaining tissue structure and function [1-6]. Residual stress in neural tissue in the brain remains incompletely characterized. Knowledge of residual stress distributions in the brain could inform predictions of brain deformation during neurosurgery, where such deformations can compromise the precision of surgical resections [7,8], and could improve models of brain tissue deformation in traumatic brain injury (TBI), which currently assume a zero residual stress state [9-12].

An additional reason to measure residual stress in the brain is to understand how mechanics contributes to brain morphology during development. Residual stress originates from processes such as tissue growth and remodeling [13]. Stresses facilitate the transmission of physical signals and regulate critical biological processes, including cell migration and axonal growth [14-22]. In humans and other mammals, the cerebral cortical surface is characterized by gyri (convex folds) and sulci (concave folds). Folding aberrations are associated with neurological conditions such as schizophrenia and autism spectrum disorder [23-26]. The mechanobiology of gyral and sulcal formation remains a topic of active research. While the influence of mechanical factors on abnormal folding patterns has been explored in theoretical and mathematical terms [27-31], measurement of residual stress in the brain remains limited [32,33]. More detailed knowledge of brain residual stress could clarify how developmental processes such as growth lead to the shape and structure of the brain at maturity.

Tissue dissection has been widely employed for measuring residual stress in other soft tissues [5,6,34,35]. This general approach involves introducing controlled incisions in the tissue to release residual stresses; the resulting deformation is measured and used to estimate the stress distribution before cutting. In the current study, we extend a technique developed for measuring solid stress in tumors [5] to obtain new measurements of local residual stresses in brain tissue. Cylindrical tissue samples extracted from the brain using a biopsy needle are [5] allowed to freely deform in artificial cerebrospinal fluid (aCSF). High-resolution magnetic resonance imaging (MRI) is then employed to capture the resulting deformation of the excised cylinder; images are analyzed using inverse finite element (FE) modeling (See Supplemental Material Videos 1 and 2) to estimate the internal stresses. MRI also enables characterization of the excised tissue by delineating grey matter and white matter regions and estimating orientations of white matter fiber bundles. This approach provides comprehensive quantitative measurements of stress magnitude, orientation, and distribution, offering valuable insights into the mechanical behavior of the mammalian brain.

## MATERIALS AND METHODS

### Animal Handling and Brain Extractions

All animal procedures were performed following the relevant guidelines and regulations outlined by the Public Health Service Policy on the Humane Care and Use of Laboratory Animals and the American Veterinary Medical Association’s Guidelines for Euthanasia of Animals. These procedures received approval from the Washington University Institutional Animal Care and Use Committee, and all experiments were conducted according to the approved guidelines and regulations. A total of 10 adult female mice (C57BL/6J strain), 3-6 months, provided by The Jackson Laboratory (Bar Harbor, ME, USA) and one female Yucatan minipig were used to acquire data for this study. Mice were deeply anesthetized with isoflurane before brain extraction, typically preserving all parts except the olfactory bulb. The Yucatan minipig was euthanized by sodium pentobarbital overdose, and the brain tissue was harvested, submerged in artificial cerebrospinal fluid (aCSF), and kept at 4°C until embedded in aCSF/agarose gel.

### aCSF and Gel Preparation

aCSF was prepared by mixing 1 mM CaCl_2_, 1 mM MgSO_4_, and 4 mM NaHCO_3_ with 1×Hanks’ Balanced Salt Solution, supplemented with 0.25 mM HEPES, 3 mM D-glucose, and 0.2% (v/v) Phenol Red. The samples were embedded in agarose gel, ensuring the gel temperature was approximately 40°C before placement. The gel consisted of low melting point agar (UltraPure™ LMP Agarose, Sigma-Aldrich) dissolved in aCSF (2.8 g agar/ 100 ml aCSF).

### Biopsy Procedure to Extract Tissue Cylinders

Before the biopsy procedure, brain tissue samples were refrigerated at 4°C for approximately 10 minutes to ensure consistent tissue extraction. A biopsy needle (1.8 mm inner diameter) with a plunger attached was mounted on a micromanipulator. The samples were positioned under the biopsy needle with adequate space for the needle to create a through hole. To obtain grey matter samples from mice, the needle was adjusted to be as close to perpendicular to the brain surface as possible. After the needle passed through the tissue and gel at the region of interest, the extracted tissue was pushed from the needle into aCSF using the air inside the attached plunger. The extracted tissue cylinder was then allowed to deform freely in aCSF for 40 minutes before being embedded into the gel. Excised samples were inspected under a dissecting microscope to ensure consistency (Fig.). In mice, after extraction of the tissue cylinder, the brain was re-imaged using the same MR sequence to identify the location of the extracted cylinder. The brain after excision was imaged in an orientation as close as possible to the orientation of the initial MRI of the intact brain. Fig. 1b depicts the steps involved in data acquisition. In the minipig, tissue cylinders were obtained from thick coronal slices of the extracted brain, Fig. 1c.

**Fig. 1:**
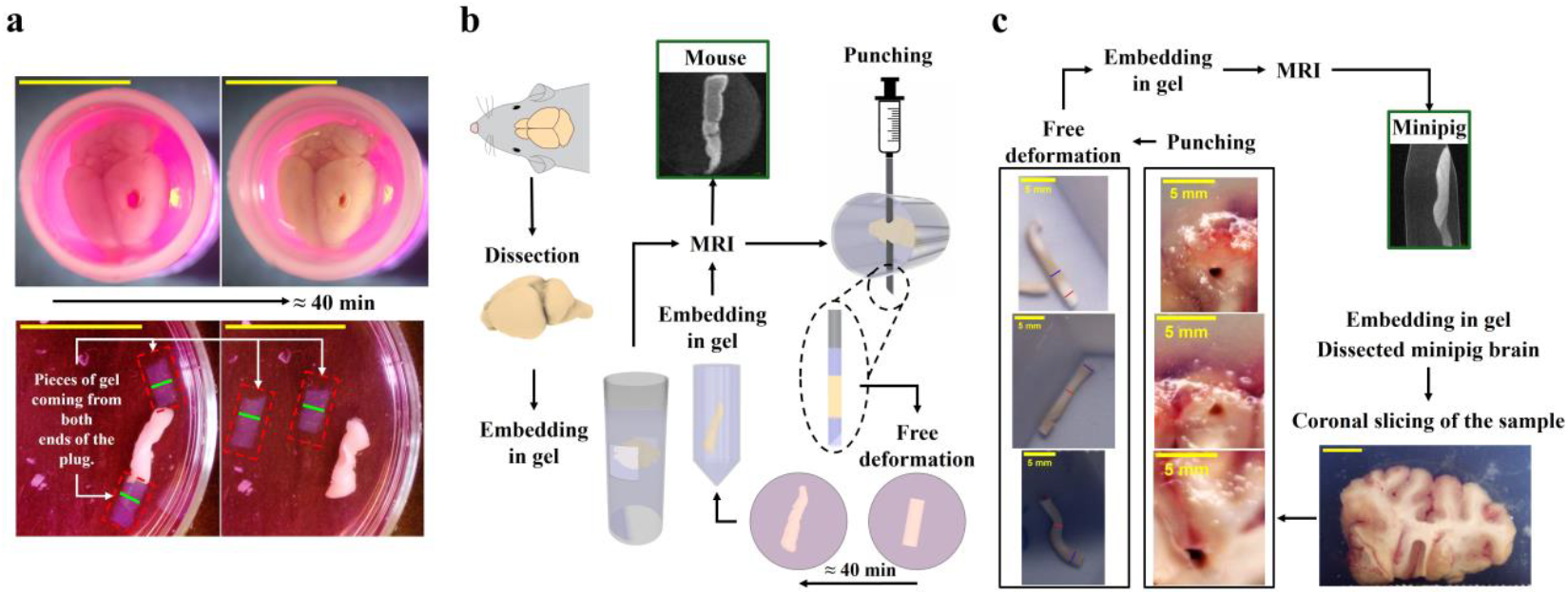
Experimental procedures. (a) Embedded mouse brain showing the hole created by tissue extraction by biopsy needle, and the corresponding excised tissue cylinder immediately after the biopsy and 40 minutes after free deformation of both the brain and the excised cylinder. Scale bars are 10 mm. (b) Experimental procedure for acquiring tissue cylinders in the mouse. (c) Experimental procedure for acquiring tissue cylinders in the minipig. Blue lines show regions of cortical grey matter; red lines indicate white matter regions (confirmed with DTI imaging). Scale bar is 10 mm in coronal slice image.

### MR Imaging of Tissue Cylinders

MR imaging was performed using a 9.4T Bruker small-animal MRI scanner. T2-weighted images of the excised tissue cylinders were acquired with an echo time (TE) of approximately 30 ms, an in-plane resolution of 75-100 μm isotropic and a slice thickness of 150-200 μm. For the minipig, in addition to T2-weighted images, diffusion tensor images (DTI) were acquired with three b-values, 200, 650, and 1000 s/mm^2^ and 20 gradient directions, to estimate the orientations of fiber tracts in the tissue cylinders. The in-plane resolution for DTI acquisition was 100 μm isotropic, and the slice thickness was 500 microns. Before post-processing DTI image volumes were resampled to an isotropic resolution of 300 microns.

### Simulation of Tissue Stress State in the Intact Brain

Simulations were performed to estimate the initial stress state in the undeformed tissue in the intact brain or tissue slice, starting from the deformed shape of the excised tissue cylinder. First, 2D sections of the MR image volume were identified in the region of interest (cortical grey matter or white matter), perpendicular to the long axis of the cylinder. In some cases, images of these axial sections of the tissue sample were defined oblique to the original imaging planes. Next, the boundary of each section was outlined by fitting a spline through manually selected points on the boundary. The initial stress state was estimated from the deformed shape of the section using commercial FE software ABAQUS™ (Dassault Systemes, Paris, France) [36]. The section was modeled with 2D hyperelastic shell elements (quadrilateral quadratic plane-stress, element type CPS8) with the maximum mesh size of 0.025 mm. To approximate stresses of the intact tissue, boundary displacements were imposed to restore the free tissue to its original circular shape. In the first step of the simulation, a pure radial displacement was applied to the boundary to align it with the circular inner diameter of the biopsy needle (1.8 mm). In the second step, points on the boundary were allowed to slide freely in the circumferential direction while maintaining the radius at 1.8 mm to reach the minimum strain energy state (See Supplemental Material Video 1 and See Supplemental Material Video 2). Mesh convergence analysis was conducted to ensure the reliability of the simulation results (Supp. Fig. 5).

The material was modeled as homogeneous neo-Hookean material with the strain energy function, Ψ, per unit of reference volume is defined as

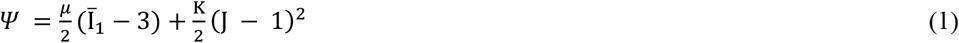

Where 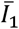 is the first deviatoric strain invariant and *J* = det ***F*** is the Jacobian of ***F, F*** is the deformation gradient tensor. We assumed a shear modulus, μ = 2 kPa, and a bulk modulus, K, 1000 times the shear modulus to approximate incompressibility. The shear modulus value was selected to approximate measured values from our own experiments [37] and reported in the literature for both mice and minipig [38-42]. Additionally, a parameter sweep on the ratio between bulk and shear moduli was performed to evaluate the effects of the incompressibility assumption on the simulation outcomes (Supp. Fig. 6).

### Statistics

We averaged the estimated absolute principal stresses (σ_1_, σ_2_) and absolute principal logarithmic strains (ϵ_1_, ϵ_2_) for cortical grey matter and white matter sections in each species (Supp. Table 1 and Supp. Table 2). A t-test [43] was used to compare the mean values of σ_1_, assessing differences in both cortical grey matter and white matter between the mouse and the minipig. A p-value of less than 0.05 was considered statistically significant for this test. To quantify the relationship between principal stress and fiber axes, the direction of maximum diffusivity, 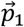 (the eigenvector corresponding to the major eigenvalue, λ_1_, of the local diffusion tensor) was compared to direction of maximum stress, 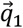 (the eigenvector corresponding to the major eigenvalue σ_1_ of the stress tensor). The angle between 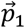, and 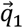 was calculated at each voxel, and significance of alignment was assessed using Rao’s spacing [44] non-uniformity test of the angle distribution [45].

## RESULTS

### Cortical Grey Matter of the Mouse Brain Exhibits Compressive Stress

Tissue columns were extracted from fresh mouse brains and imaged with MRI to obtain the deformed 3D geometry of the relaxed tissue (Fig. 2a). In the mouse, tissue columns were extracted with the biopsy needle oriented perpendicular to the cortical surface. Two-dimensional (2D) cross-sections in planes parallel to the cortical surface, within the cortical grey matter (Fig. 2b), were analyzed by inverse FE simulation (See Supplemental Material Video 1). These sections consistently exhibited compressive in-plane stress, as evident from the expansion from the original 1.8 mm diameter size (green circle, Fig. 2b) to a larger diameter in the relaxed tissue (red contour, Fig. 2b).

**Fig. 2:**
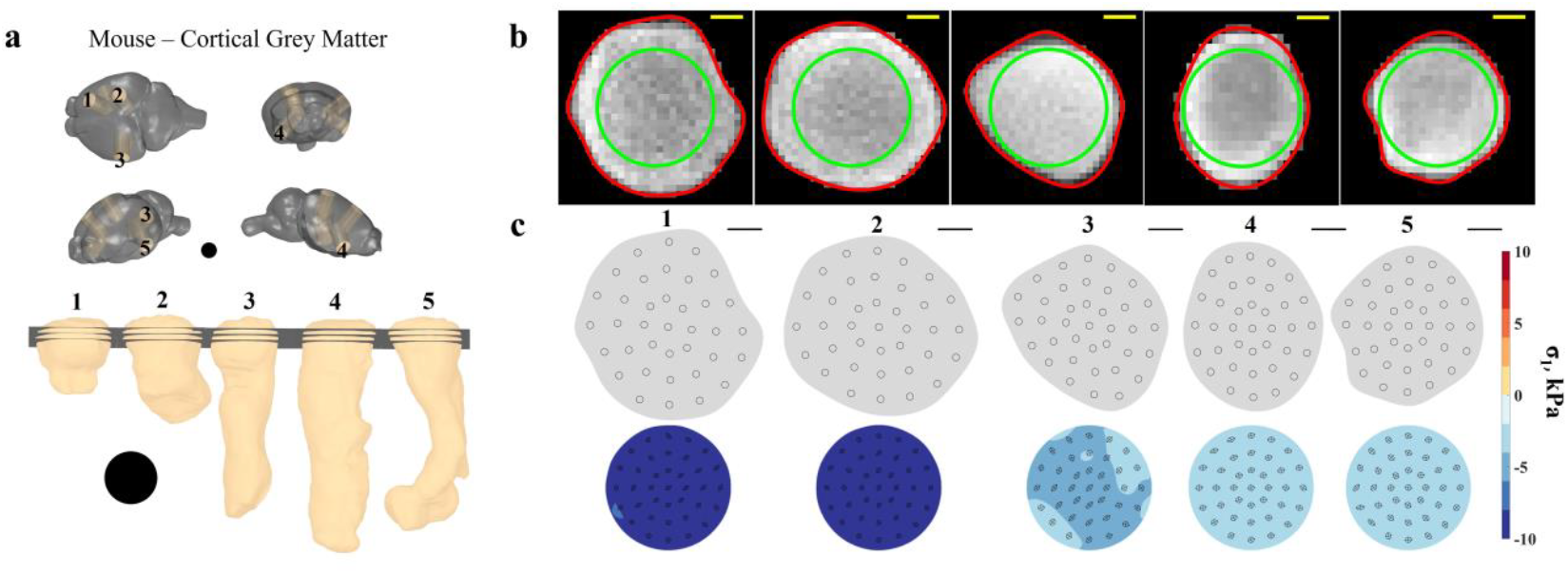
Residual stress in cortical grey matter of the mouse brain. (a) Locations of tissue sample extraction and sections from the cylindrical samples. To acquire these samples, the biopsy needle was oriented perpendicular to the cortical surface. Tissue volumes are reconstructed from MR images, black circles depict the inner diameter of the biopsy needle (1.8 mm). (b) MR images of the middle section for each trio of analyzed sections; green circles indicate the size of the biopsy needle, and the red curves outline the relaxed, deformed shape used for stress estimation. (c) Top row: section shapes created from MR image slices of excised cylinders of cortical grey matter. Bottom row: simulation-predicted stress fields in each section prior to relaxation (i.e., stress fields generated by elastic deformations to original circular shape). Color bar indicates the maximum absolute principal stress (σ_1_). Ellipses depict the local deformation of circles with an initial radius of 0.05 mm in the relaxed state. Scale bars represent 0.5 mm.

Estimated stresses in sections containing only grey matter are shown in Fig. 2c. Mean values for both in-plane principal stress components (σ_1_ and σ_2_) were found to be negative (compressive) for all sections. (Magnitudes of principal stresses and strains are further described in supplemental material; Supp. Table 1.) The state of compression was observed within all cortical grey matter sections analyzed (Fig. 2).

### White Matter in the Mouse Brain Exhibits Tensile Stress

To characterize residual stresses in white matter, sections of tissue columns beneath the cortical layer were also analyzed. At a depth of approximately 1 mm below the cortical surface, the intensity of T_2_-weighted MR images was reduced in all or part of the tissue column, indicating the presence of myelinated white matter (Fig. 3). Within these image slices, tissue was observed to contract following extraction from the brain; red contours encircle an area that is smaller than the original needle bore, indicated by green circles (Fig. 3b; See Supplemental Material Video 2). Thus, in contrast to observations of the cortical grey matter, tissue sections from the mouse that contain white matter exhibited in-plane tension. Notably, the observed tension was anisotropic and the direction of maximum tension tended to align with the direction of the band of white matter.

**Fig. 3:**
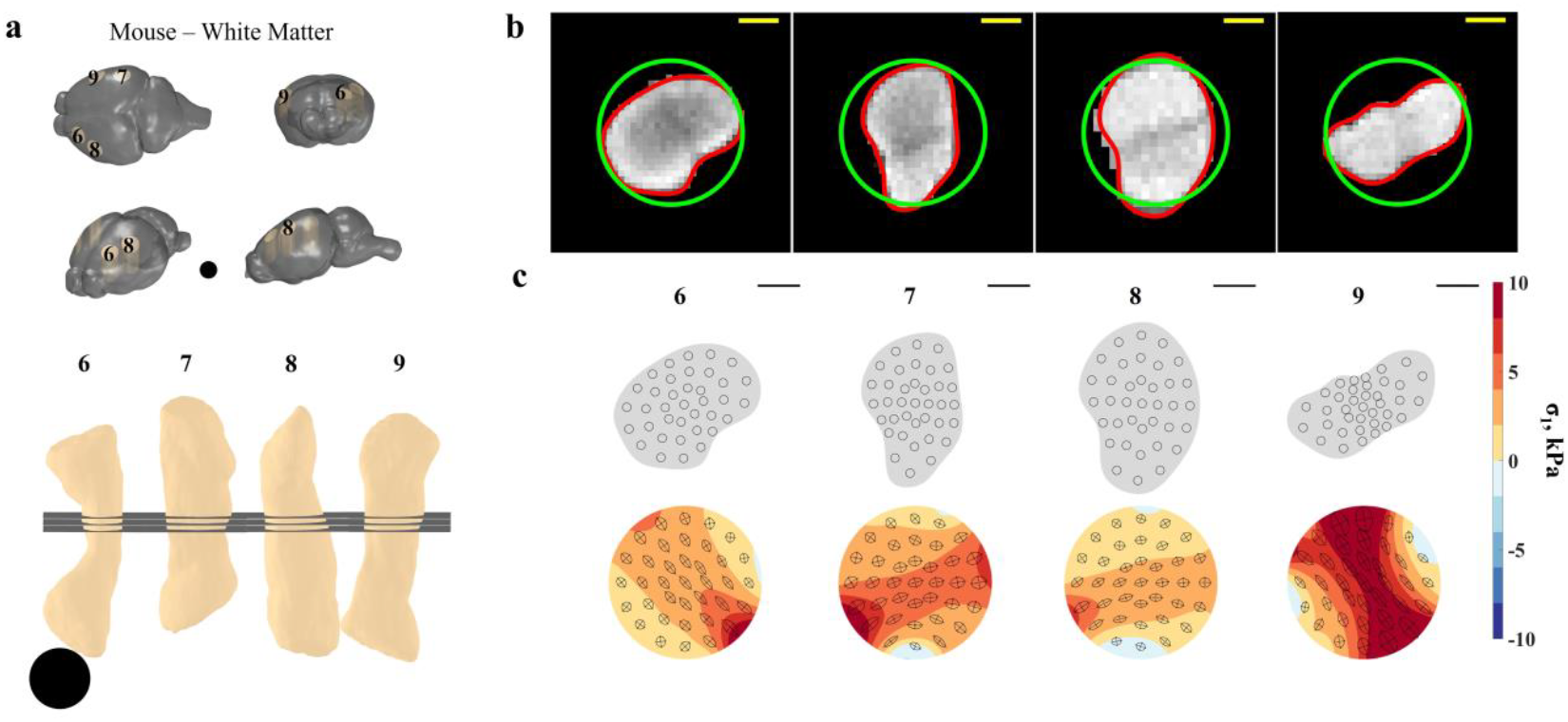
Residual stress in white matter of the mouse brain. (a) Locations of tissue extraction and sections from the extracted samples. Tissue volumes are reconstructed from MR images, black circles depict the inner diameter of the biopsy needle (1.8 mm). (b) MR images of the middle sections for each sample containing white matter tracts; green circles indicate the inner diameter of the biopsy needle, and the red curves outline the relaxed, deformed shape used for simulation. (c) Top row: section shapes created from the middle section of the trio of analyzed sections containing white matter. Bottom row: simulation-predicted stress fields in each section prior to relaxation (i.e., stress fields generated by elastic deformations to original circular shape). Color bar indicates the maximum absolute principal stress (σ_1_). Ellipses represent the local deformation of circles with an initial radius of 0.05 mm in the relaxed state. Scale bars represent 0.5 mm.

Maps of maximum absolute principal stress (σ_1_) estimates in sections of mouse brain tissue containing white matter are shown in Fig. 3c. Average values for both σ_1_ and σ_2_ were found to be positive (tensile) in these sections (Fig. 3b); corresponding strain estimates are included in supplementary material; SeeSupp. Table 2. These findings indicate that tension is the dominant feature of residual stress in murine white matter and suggest that tension is largest along the dominant axis of the fiber bundle.

### Stress in Grey Matter and White Matter of a Gyrencephalic (Minipig) Brain

To investigate if differences in stress between grey and white matter were also present in a larger gyrencephalic mammal, tissue columns were obtained from the brain of Yucatan minipig. In contrast to the mouse, white matter structures in the minipig are large relative to the 1.8-mm diameter of the biopsy needle (Fig. 4a). Maps of maximum (absolute) principal stress σ_1_ in sections of cortical grey matter from the minipig brain (Fig. 4b) in gyri (samples 10-11) and sulci (sample 12) exhibited bi-axial in-plane compression similar to cortical samples obtained from mouse brains (See Supp. Table **1**). Maps of estimated maximum absolute principal stress, σ_1_, in white matter sections from the minipig brain are shown in Fig. 4c. Regions containing purely white matter in the minipig brain exhibited deformations consistent with anisotropic tension (Fig. 4c). The magnitude of maximum tension was similar for white matter under sulcal and gyral regions (See Supp. Table **2**). As in mouse brain sections that contain white matter, anisotropy of tension is evident from the direction-dependent shrinkage of minipig brain sections containing white matter.

**Fig. 4:**
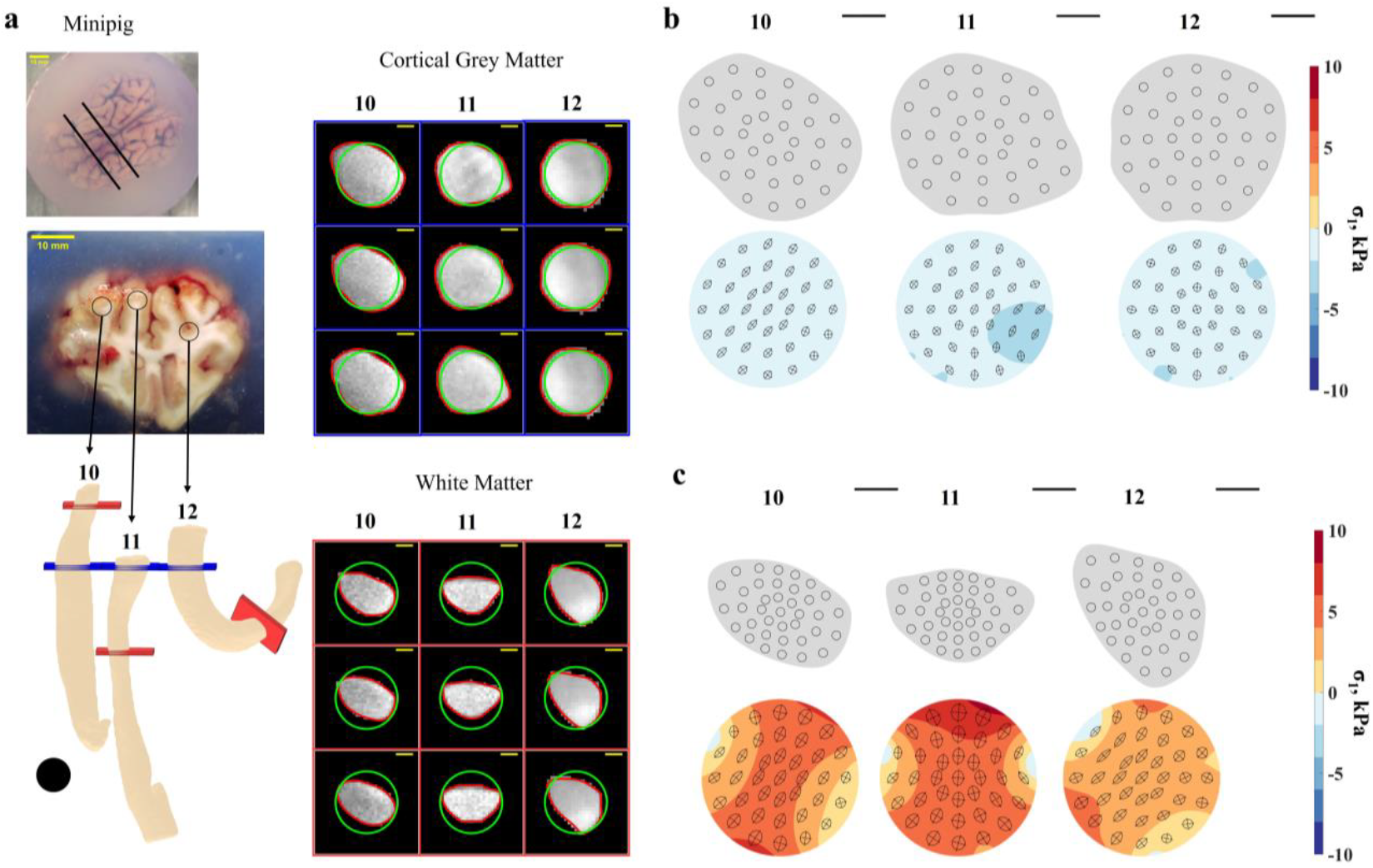
Residual stress in cortical grey matter and white matter in the minipig brain. (a) Locations of acquired tissue samples and sections from the samples. In the minipig the biopsy needle was oriented perpendicular to the coronal slab, roughly parallel to the cortical surface. Tissue volumes are reconstructed from MR images; black circles represent the inner diameter of the biopsy needle (1.8 mm). MR images of tissue sections (right) depict original biopsy needle diameter (green circle) and relaxed shape (red outline). (b) Maps of estimated maximum absolute principal stress, σ_1_, in sections of excised cylinders of minipig brain cortical grey matter. Top row: shapes of sections from minipig samples containing mainly cortical grey matter. Bottom row: simulation-predicted stress fields in the corresponding sections prior to relaxation. (c) Maps of estimated maximum (absolute) principal stress, σ_1_, in in sections of excised cylinders of minipig brain tissue containing mostly white matter. Top row: shapes of sections from minipig samples containing white matter. Bottom row: simulation-predicted stress fields in the corresponding sections prior to relaxation. Scale bars represent 0.5 mm. Color bar indicates the computed σ_1_ in corresponding section. Ellipses represent the local deformation of circles with an initial radius of 0.05 mm in the relaxed state.

### The Direction of Tension Aligns with the Direction of White Matter Tracts

Reconstructed images of the mouse brain show approximately isotropic compressive in-plane stress in the cortical gray matter (Fig. 2). In contrast, mouse brain tissue sections containing white matter (Fig. 3), the stress is anisotropic and tensile and the maximum tension direction generally aligns with the white matter tracts.

To directly investigate the relationship between the direction of tension and the orientation of axon fibers, water diffusion anisotropy was measured within tissue column slices of white matter from the minipig brain. Sections of white matter had higher fractional anisotropy (FA) of water diffusion than sections of grey matter within the same tissue column (Fig. 5a); this is expected since white matter is made up of aligned, myelinated axonal fibers that restrict radial diffusion relative to axial diffusion. To indicate the direction of the underlying white matter axon fiber bundles, the direction of maximum diffusivity (the primary eigenvector of the diffusion tensor) 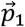, was projected onto the image plane at each voxel (Fig. 5a, blue lines). The 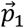 were generally aligned with the direction of maximum tensile stress (the primary eigenvector of the 2D stress tensor), 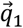, mapped onto the deformed section, indicated with dashed red lines in Fig. 5a. The distribution of angles between the directions of maximum stress (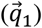) and maximum diffusivity 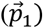 in white matter sections is shown in Fig. 5b; the correlation between the two directions is statistically significant (p<0.001; Rao’s spacing non-uniformity test [44]).

**Fig. 5:**
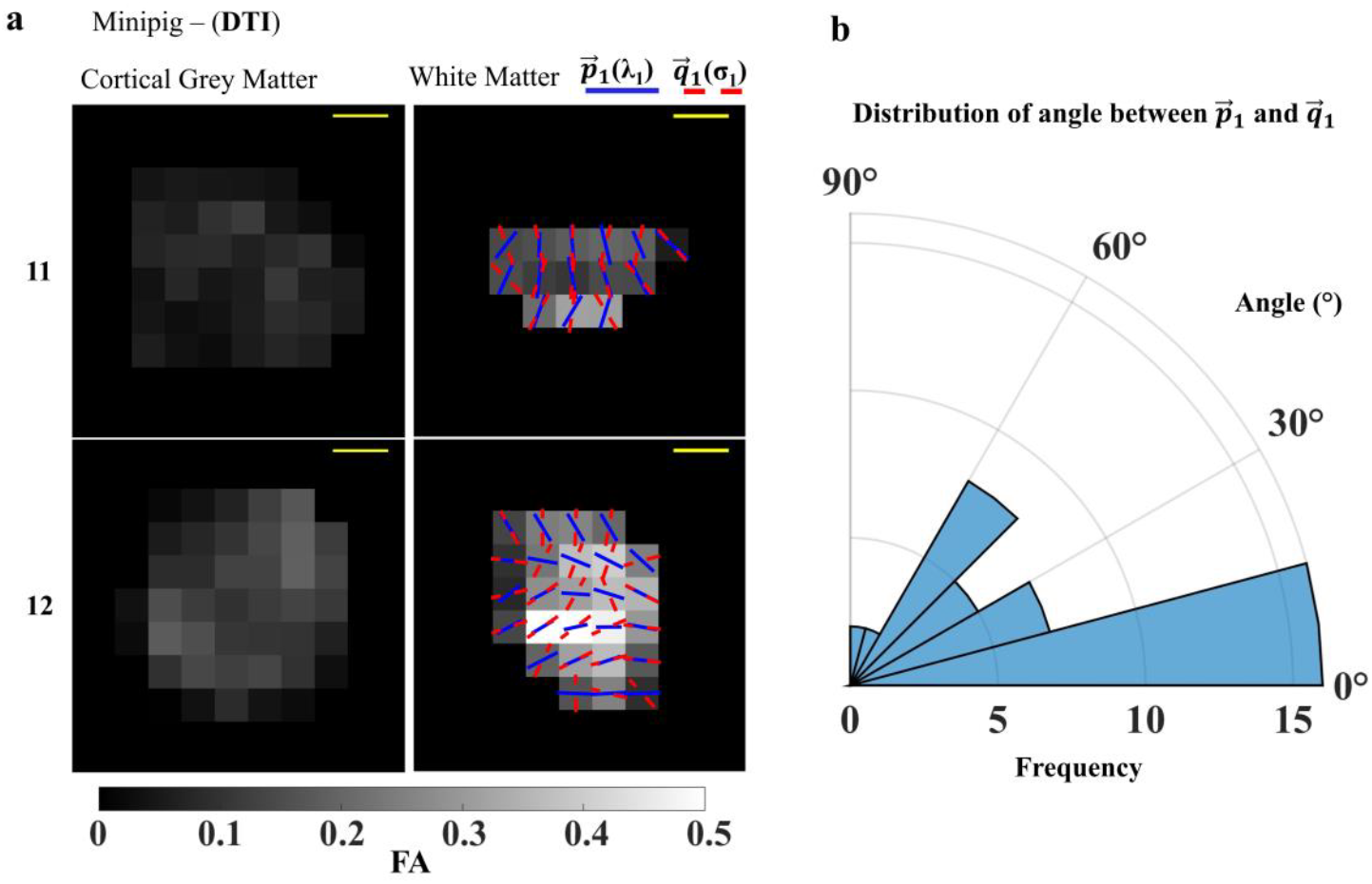
Comparison of anisotropic diffusion and tension in minipig white matter. (a) Diffusion tensor imaging results from the minipig brain (samples 11 and 12). Maps of fractional anisotropy (FA) for cortical grey matter sections (left) and white matter sections (right). Solid blue lines show the in-plane projection of the direction of maximal diffusivity 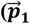, ∼ fiber orientation) and dashed red lines show the direction of maximum in-plane tension 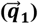. (b) Distribution of angle between 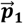 and 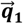 in both samples sowing correlation between tension and fiber axes (p< 0.001).

### Summary of Residual Stress in Mouse and Minipig Brains

In both the mouse and minipig, grey matter is under compressive stress whereas white matter is in a state of anisotropic tension, as summarized for all samples in Fig. 6. In samples from the mouse, the compressive stress in cortical gray matter appears to vary with cortical region. The magnitude of compressive stress generally appeared to be higher in samples from the mouse cortex than from the minipig cortex, however in the mouse the stress was estimated in a plane parallel to the brain surface but in the minipig the planes of analyzed sections were not parallel to the cortical surface.

**Fig. 6:**
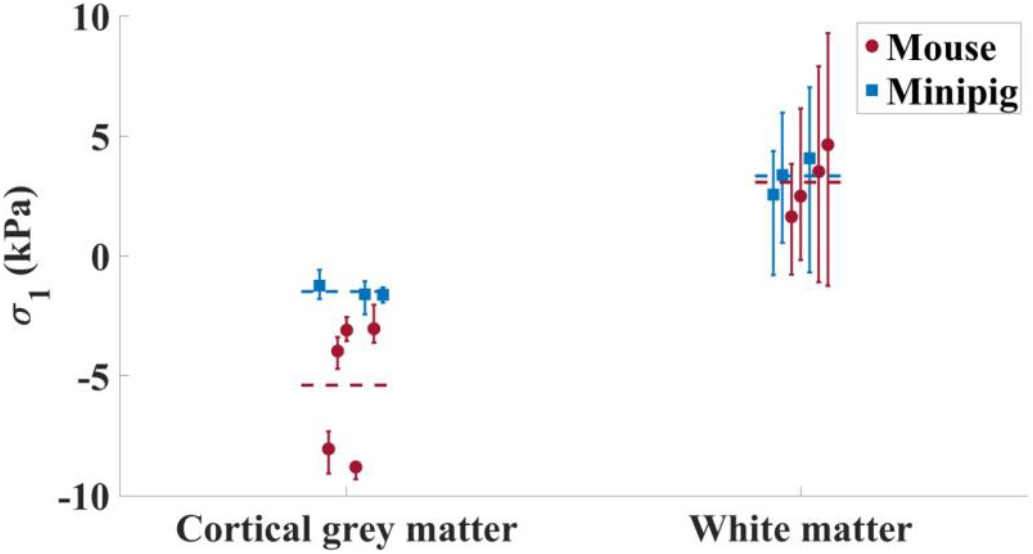
Average values of maximum (absolute) principal stress, σ_1_, in each tissue sample. Error bars indicate the 95% confidence interval of the estimated values. Dashed lines depict the average value of stress for samples from mouse (red) and minipig (blue).

## DISCUSSION

We quantified residual stress in grey and white matter brain tissue in a lissencephalic rodent (the adult mouse) and a gyrencephalic large mammal (the minipig) by extracting cylindrical tissue samples, obtaining high resolution MR images of the relaxed, deformed tissue shape, and performing inverse finite element simulations of the deformation process. In prior studies, optical coherence tomography images of tissue cylinders extracted by biopsy needle have been analyzed to estimate solid stress in tumors[5]. The current study extends this approach, using images from high-resolution MRI to illuminate the stress state in the brain. The current findings confirm previous, more limited studies based on simple incisions that indicated compression in cortical grey matter and tension in white matter. By measuring 2D deformation of sections of extracted tissue cylinders we are able to obtain quantitative estimates of the 2D stress state in the tissue.

### Implications for Brain Development

Xu et al. [32] made incisions in coronal slices of the adult mouse brain to investigate the stress state in the mature rodent brain. Cuts perpendicular to axonal fiber tracts caused the tissue to open, while cuts in gray matter remained closed. These results indicated that the cortical grey matter is compressed (or possibly under zero stress), and the white matter is in tension. Subsequent experiments in the larger, gyrencephalic ferret brain (both adult and juvenile) [33] generated similar results, with the additional observation that cuts in the white matter opened only in cuts perpendicular to the fiber axis; opening occurred in the direction of fiber alignment. However, cuts only reveal the sign of internal forces perpendicular to the incision and, in the case of a cut that does not open, cannot distinguish between high compression, low compression, or even a zero-stress state.

In contrast, the current approach interrogates the complete in-plane (2D) stress state in a section of tissue, providing maps of the magnitude and direction of the maximum absolute principal stress within the different tissue regions. In the mouse, estimates of compressive stress in the cortical layer, Fig. 2, and tensile stress in the white matter tract (Fig. 3, Fig. 5) are consistent with behavior reported by Xu et al. [32] and confirm the contrasting stress state of cortical grey matter and white matter. Going beyond prior observations, we were able to estimate the magnitudes of compression within the cortical grey matter and tension in white matter in the mouse. We determined that tangential compression within the cortex is relatively isotropic (similar values for both principal stress values in planes parallel to the cortical surface).

In the minipig brain, we measured stress states consistent with the more qualitative results of previous incision experiments of Xu et al. [33] in the adult ferret brain. As in the mouse and ferret, cortical grey matter in the minipig is under compression (Fig. 4b), and subcortical white matter is under tension (Fig. 4c). The current study provides new information quantifying the magnitude of stress and confirms that the direction of maximum tension in minipig white matter (Fig. 5c) approximates the fiber orientation (as estimated from the direction of maximum water diffusivity). This observation complements recent simulations of cortical expansion-driven folding that have illustrated that mechanical stress may help guide or orient fibers in white matter [46]. In the convoluted brains of higher mammals, the fiber axis is predominantly radial under gyri and circumferential under sulci [47,48]. The distribution of residual stress and the alignment of the maximum tension direction with fiber direction in the minipig brain agree with the predictions of cortical expansion-driven models of folding in which the growth of axons is mediated by stress [46].

Substantial regional variation in compressive stress was observed in the mouse cortical grey matter (Fig. 2, Fig. 6), with compression apparently higher in samples from dorsal than lateral cortex. These regions have been previously shown to exhibit increased cortical thickness [49,50]. Thus, our results tentatively suggest that thicker cortical regions experience higher compressive stress. Rajagopalan et al. [51] observed localized cortical thickening in the fetal human brain before fold formation, while Costa Campos et al. [52] highlighted the role of cortical thickness in folding patterns. These observations are consistent with the postulate that cortical regions that experience compressive stress will first thicken, ultimately buckling if that stress exceeds a critical value. The estimated stress in the convoluted minipig cortex (Fig. 4b, Fig. 6) is consistent with the idea that folding arises from a buckling-like instability, with the observed lower stress possibly resulting from the stress release after buckling.

Several studies have measured axonal elongation and tensile stress limits of axons [18-21], leading to predictions of sustained tension in white matter ranging between 0.1-10 kPa [32]. Our estimates of tension in white matter (Fig. 6) fall within this range. The similarity in estimated tension between the lissencephalic mouse brain and gyrencephalic minipig brain suggests the possibility of a universal target stress of on the order of 2-4 kPa in white matter, beyond which white matter grows to relieve excess tension. The similarity of tensile stress values in white matter in samples from different regions also tends to support the concept of a general target stress for axons [27].

### Implications for Brain Injury and Neurosurgical Interventions

Our observations of different stress states in grey and white matter have implications not only for brain development, but also TBI, neurosurgery and neurotechnology. In the context of TBI, residual stresses in the brain’s natural equilibrium state – or preloading – will affect the brain’s response to applied external loads. Compressive stress in grey matter may confer additional stiffness [22], while tensile stress in white matter may enhance the susceptibility of axons to elongation and damage during impacts [53,54]. Incorporating residual stress in computational models of TBI mechanics could improve predictions of injury and strategies for prevention [55,56].

Neurological interventions, such as surgery or drug delivery, are also affected by residual stress. Tissue stress affects brain behavior after surgical incisions or device insertion, including differential deformations in grey and white matter [57-63], and affect the precision of resections or drug delivery [64,65].

A limitation of this study is that measurements are performed in tissue ex vivo and in vitro. Despite careful handling, including submersion in aCSF or embedding in aCSF/agarose gel to minimize drying or swelling, the material properties of brain tissue inevitably differ after death. Brain extraction and manipulation of tissue may also introduce measurement artifacts; we attempt to mitigate these effects by careful handling and allowing tissue to equilibrate in aCSF before imaging. We also note that the 2D stress tensor estimated from the deformation of a planar tissue section does not fully capture the complexity of stress distribution within the brain. Estimates of stress rely on a simplified material model using parameters from our prior studies [37]. Generalizing the current approach to 3D and developing methods to estimate tissue stress in vivo are challenging but compelling topics of future investigation.

## DATA AVAILIBILITY

Data will be shared upon request.

## ACKNOWLEDGEMENT

We acknowledge financial support from National Institutes of Health grants R01 NS111948.

## AUTHOR CONTRIBUTIONS

R.B. performed the experiments, analyzed the data, created figures, and wrote the manuscript. C.D.K., K.E.G., and P.V.B. conceived the study, supervised the experimental procedures and data analysis, and contributed to manuscript editing.

## SUPPLEMENTAL MATERIAL

**Supp. Fig. 1:**
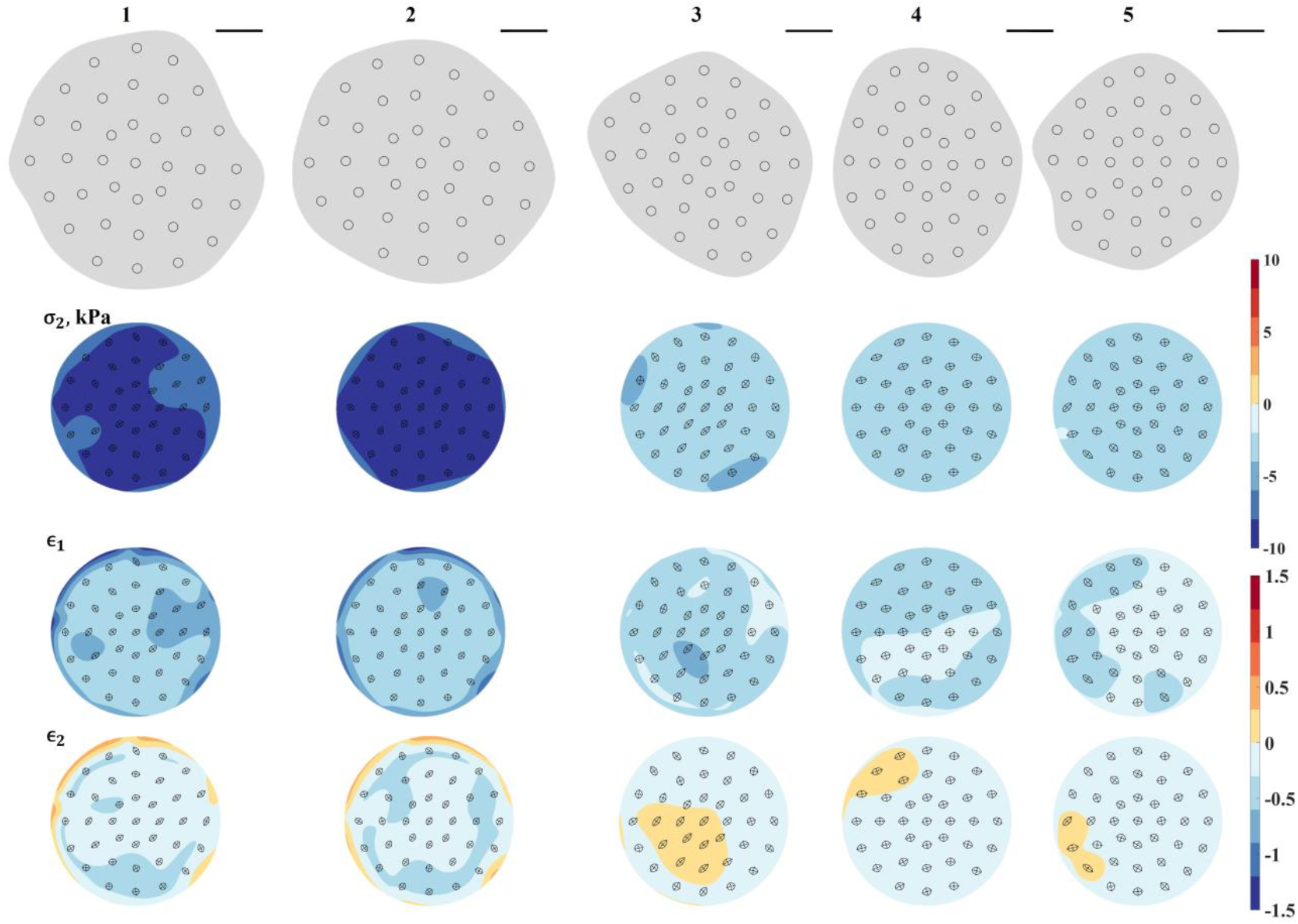
Simulation-predicted maps of minimum absolute principal stress, σ_2_, maximum absolute principal logarithmic strain, ϵ_1_, and minimum absolute principal logarithmic strain, ϵ_2_, for sections 1-5 obtained from mouse grey matter.

**Supp. Fig. 2:**
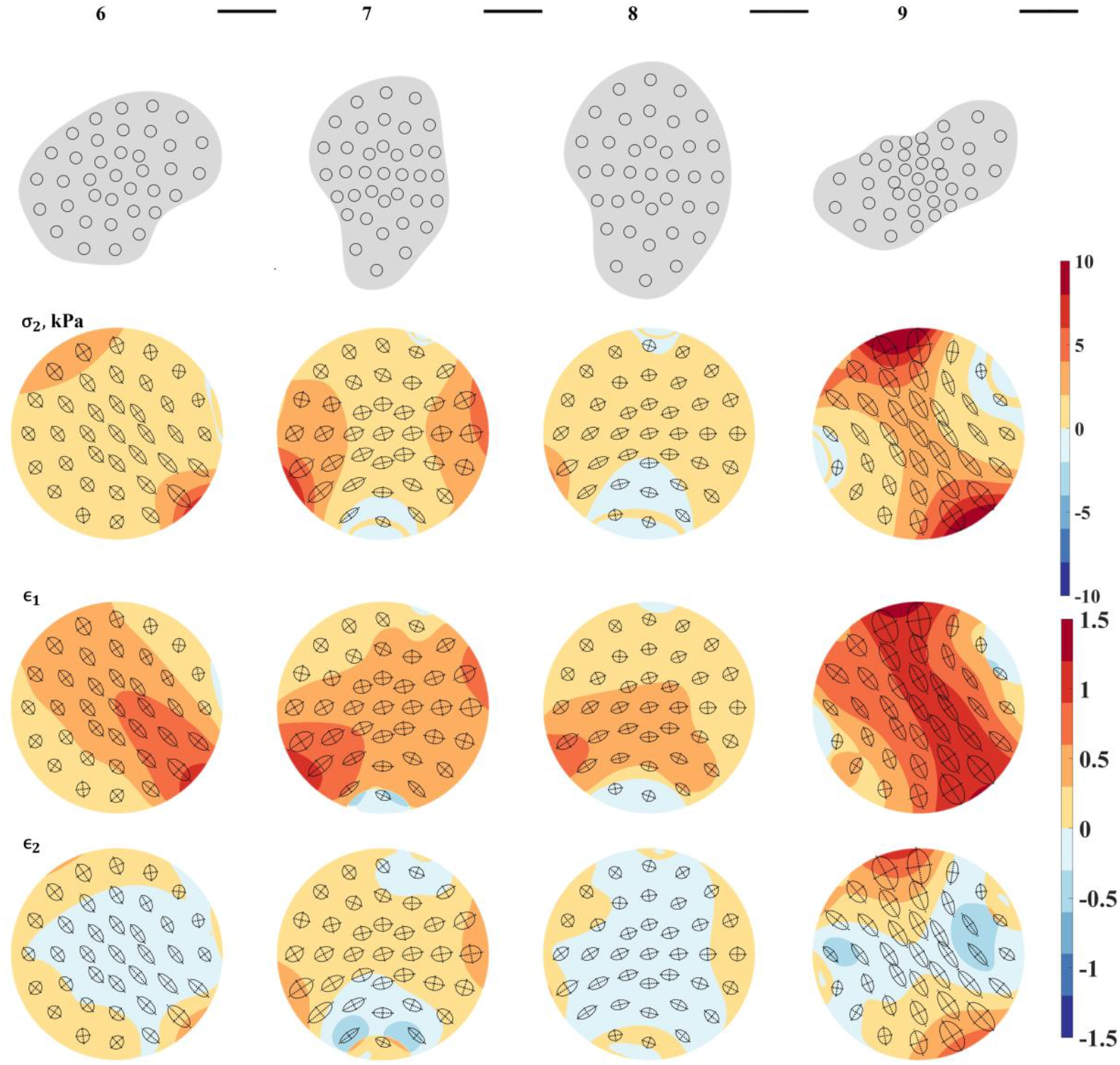
Simulation-predicted maps of minimum absolute principal stress, σ_2_, maximum absolute principal logarithmic strain, ϵ_1_, and minimum absolute principal logarithmic strain, ϵ_2_, for sections 6-9 obtained from mouse white matter.

**Supp. Fig. 3:**
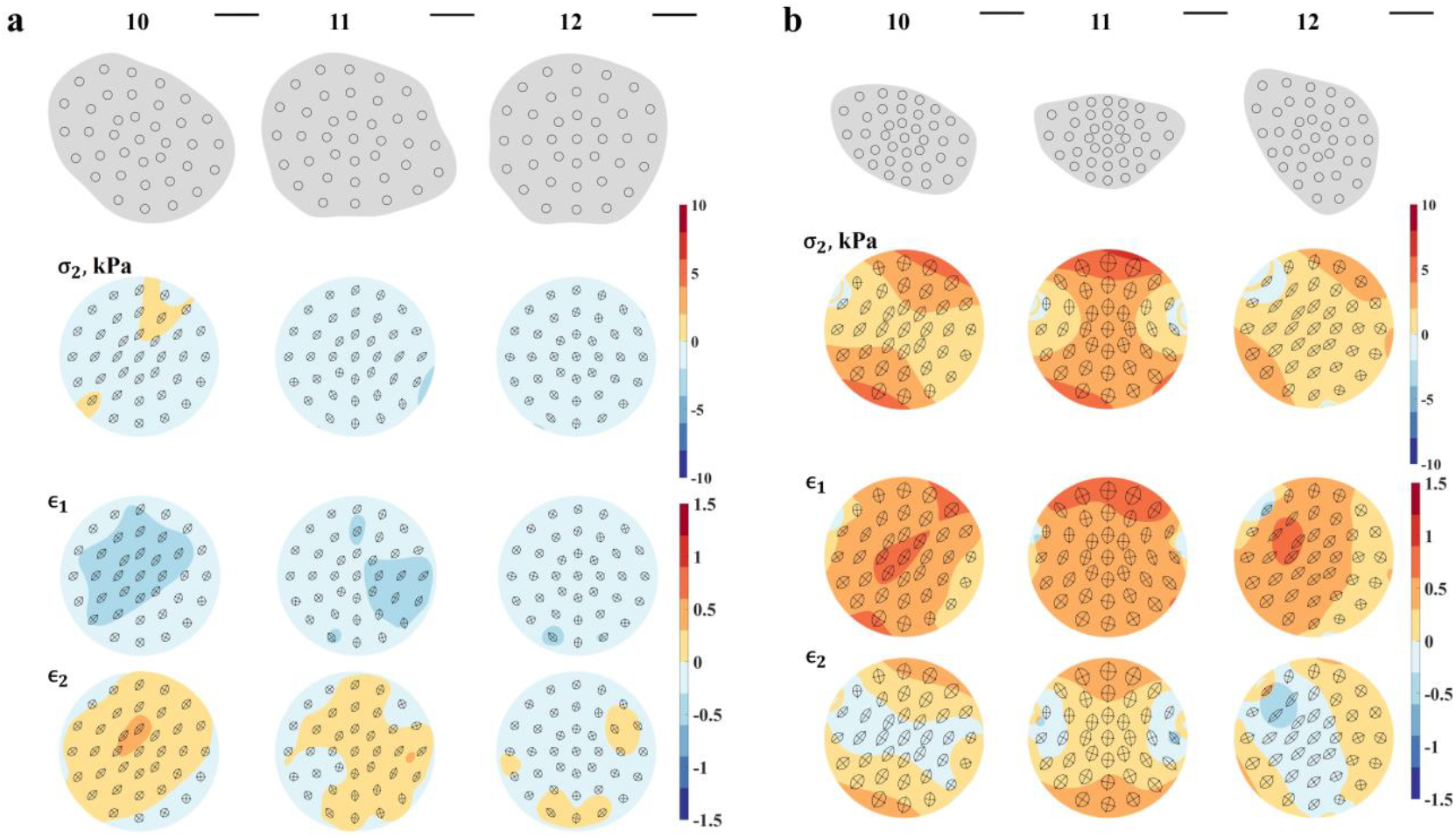
Simulation-predicted maps of minimum absolute principal stress, σ_2_, maximum absolute principal logarithmic strain, ϵ_1_, and minimum absolute principal logarithmic strain, ϵ_2_, for samples 10-12 in (a) minipig grey matter and (b) minipig white matter.

**Supp. Fig. 4:**
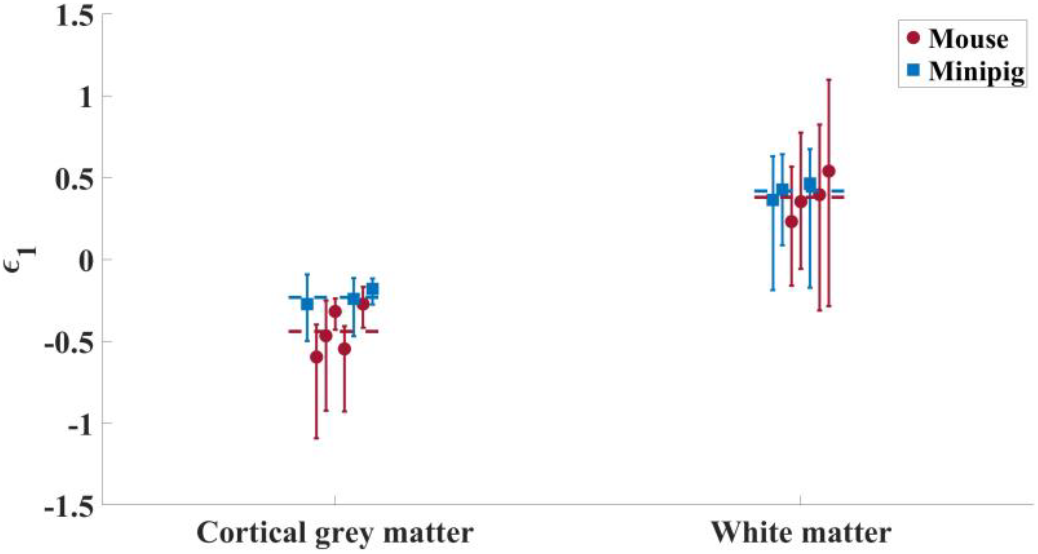
Average values of maximum (absolute) principal strain, ϵ_1_, within each sample. Error bars indicate 95% confidence interval of the computed values. Dashed lines are averages over all samples for each species.

**Supp. Fig. 5:**
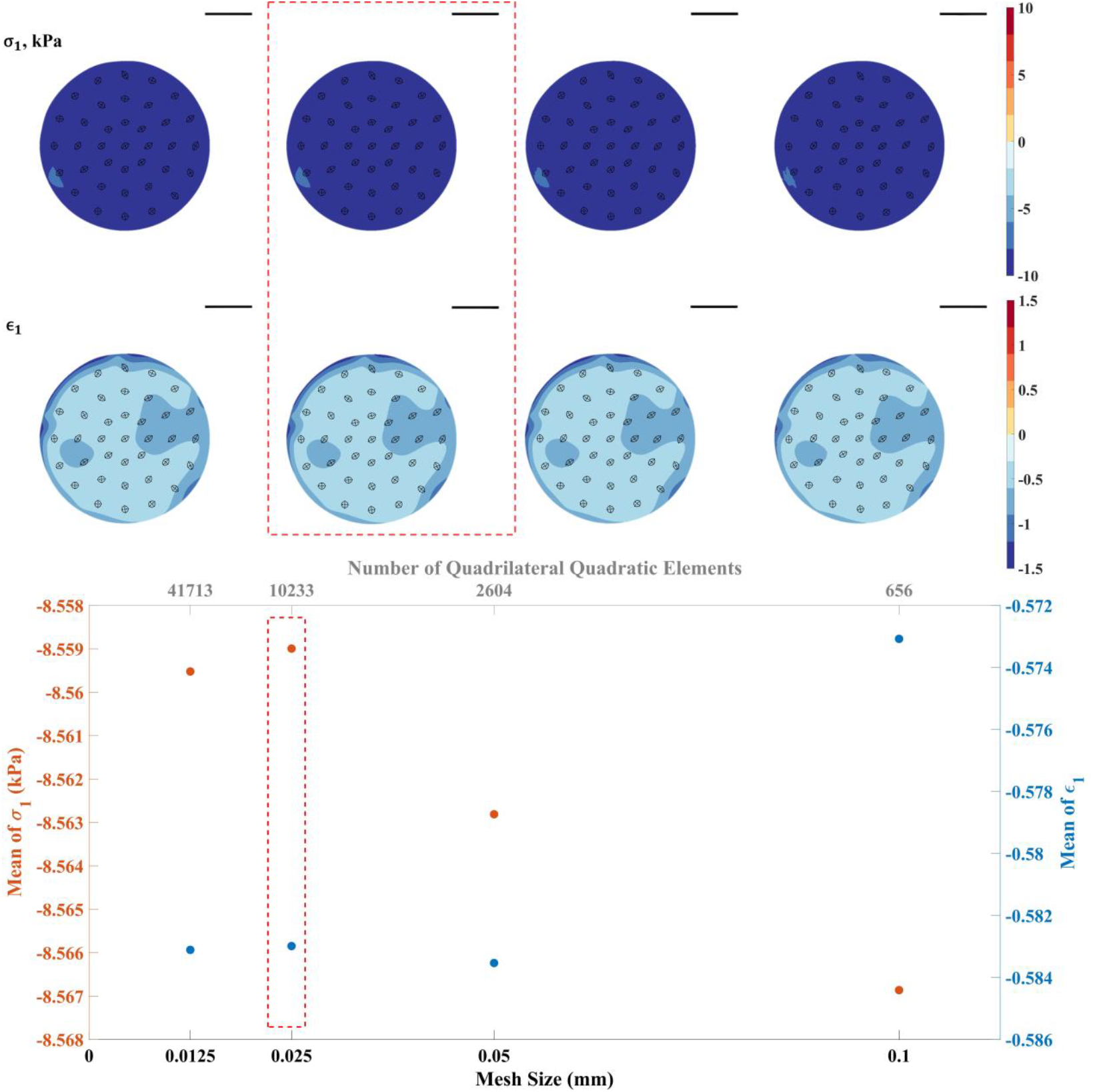
Mesh convergence study for the simulation-predicted stress field in a cortical brain tissue section from the mouse (sample 1 in the manuscript).

**Supp. Fig. 6:**
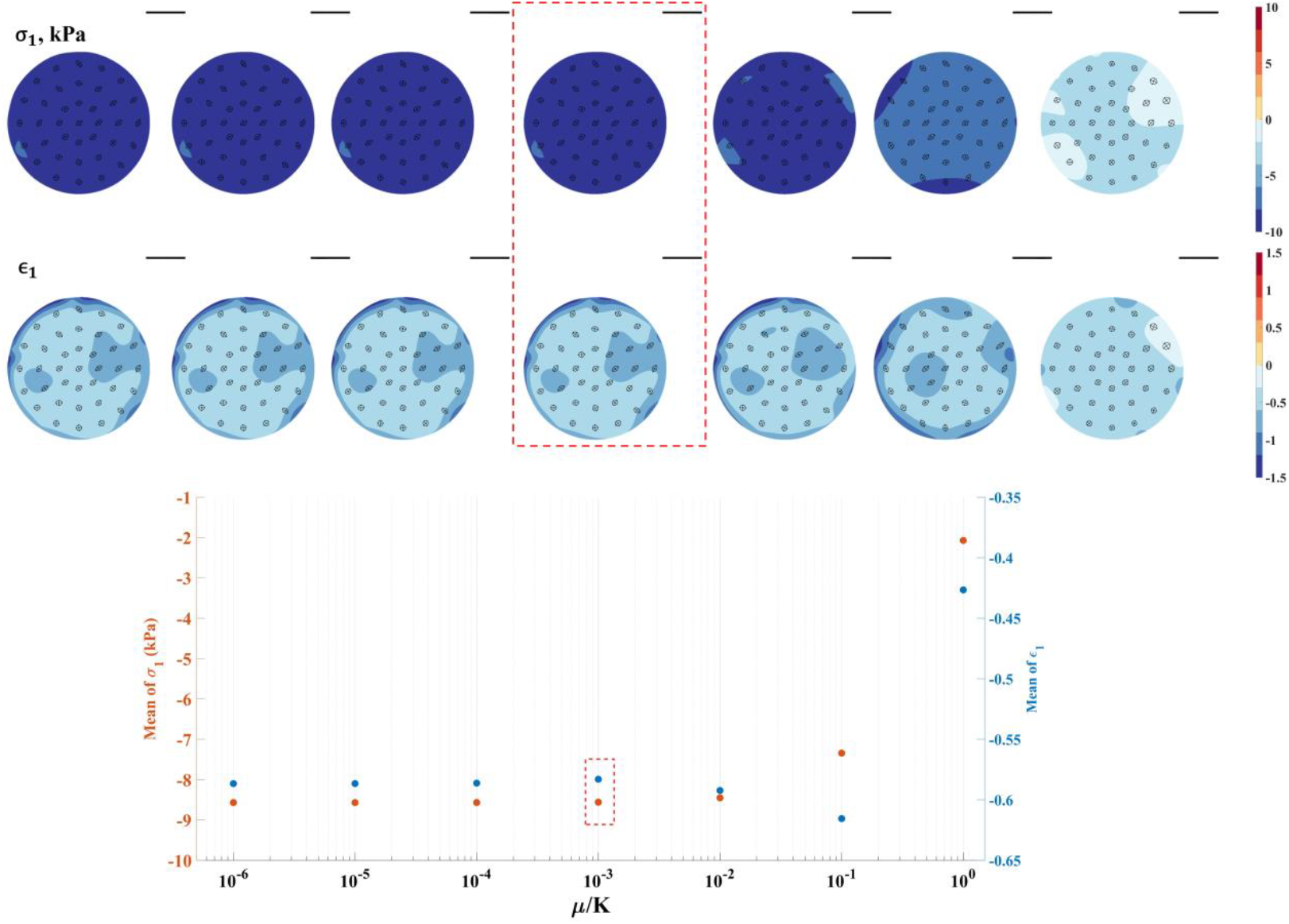
Effect of the assumed ratio of shear modulus to bulk modulus on simulation-predicted stress fields in a section of cortical brain tissue in the mouse (sample 1 in the manuscript).

**Supp. Table 1:** Averaged stress and strain estimates for grey matter samples

| Sample<br>Number | 1 | 2 | 3 | 4 | 5 | 10 | 11 | 12 |
| --- | --- | --- | --- | --- | --- | --- | --- | --- |
| | Mean ± Std | | | | | | | |
| $\sigma_1$ (kPa) | -8.05±0.45 | -8.61±0.28 | -3.97±0.16 | -3.10±0.34 | -3.00±0.61 | -1.24±0.04 | -1.61±0.18 | -1.62±0.04 |
| $\epsilon_1$ | -0.60±0.04 | -0.55±0.02 | -0.47±0.11 | -0.32±0.01 | -0.27±0.03 | -0.27±0.02 | -0.24±0.01 | -0.18±0.01 |
| $\sigma_2$ (kPa) | -7.41±0.59 | -8.21±0.31 | -3.23±0.46 | -2.45±0.36 | -2.70±0.66 | -0.43±0.07 | -0.98±0.21 | -1.31±0.03 |
| $\epsilon_2$ | -0.15±0.06 | -0.23±0.03 | 0.02±0.12 | -0.05±0.02 | -0.10±0.04 | 0.14±0.02 | 0.04±0.02 | -0.04±0.01 |

**Supp. Table 2:** Averaged stress and strain estimates for white matter samples

| Sample<br>Number | 6 | 7 | 8 | 9 | 10 | 11 | 12 |
| --- | --- | --- | --- | --- | --- | --- | --- |
| | Mean ± Std | | | | | | |
| $\sigma_1$ (kPa) | 2.49±0.89 | 3.55±1.01 | 1.63±0.16 | 4.64±0.12 | 3.37±0.30 | 4.07±0.22 | 2.56±0.27 |
| $\epsilon_1$ | 0.35±0.09 | 0.40±0.1 | 0.23±0.01 | 0.54±0.01 | 0.43±0.03 | 0.46±0.03 | 0.37±0.04 |
| $\sigma_2$ (kPa) | 1.13±0.57 | 1.77±0.43 | 0.49±0.13 | 1.50±0.04 | 1.76±0.17 | 2.32±0.14 | 1.21±0.14 |
| $\epsilon_2$ | -0.03±0.05 | 0.08±0.03 | -0.05±0.02 | -0.04±0.01 | 0.05±0.02 | 0.14±0.01 | -0.02±0.02 |

